# An Orthogonal T7 Replisome for Continuous Hypermutation and Accelerated Evolution in *E. coli*

**DOI:** 10.1101/2024.07.25.605042

**Authors:** Christian S. Diercks, Philipp J. Sondermann, Cynthia Rong, David A. Dik, Thomas G. Gillis, Yahui Ban, Peter G. Schultz

## Abstract

Systems that perform continuous hypermutation of designated genes without compromising the integrity of the host genome can dramatically accelerate the evolution of new or enhanced protein functions. We describe an orthogonal DNA replication system in *E. coli* based on the controlled expression of the replisome of bacteriophage T7. The system replicates circular plasmids that enable high transformation efficiencies and seamless integration into standard molecular biology workflows. Engineering of T7 DNA polymerase yielded variant proteins with mutation rates of 1.7 × 10^−5^ substitutions per base *in vivo* – 100,000-fold above the genomic mutation rate. Continuous evolution using the mutagenic T7 replisome was demonstrated by expanding the substrate scope of TEM-1 β-lactamase and increase activity 1,000-fold against clinically relevant monobactam and cephalosporin antibiotics in less than one week.

## Introduction

Low organismal mutation rates (∼10^−10^ substitutions per base pair, spb) render mere passaging of cells under selective pressure an inefficient strategy for the diversification of genes on laboratory timescales.(*1, 2*) Historically, directed evolution addressed this limitation by mutagenesis *in vitro* (e.g., error-prone PCR) followed by transformation of a microbial host with the diversified libraries for selection of phenotypic fitness.(*3*–*5*) However, such *in vitro* mutagenesis renders directed evolution time consuming and laborious and limits both the depth (rounds of mutagenesis) and scale (number of parallel experiments) of these experiments. While alternative randomization strategies (e.g., DNA shuffling) have improved the efficiency of this process, the removal of *ex vivo* diversification altogether by increasing *in vivo* mutagenicity enables continuous evolution at an accelerated pace.(*6*)

Early efforts to increase *in vivo* mutation rates relied on chemical mutagens, engineered host DNA polymerases, and impairing of host DNA repair machinery.(*7*–*9*) However, the mutation rates that can be achieved according to these strategies are fundamentally limited by the large number of essential host genes that must be maintained faithfully to ensure survival. Efforts to target mutagenesis to a defined locus or gene using translational fusions of deaminases to nCas9 or T7 RNA polymerase, as well as of nCas9 to error-prone DNA polymerases have been investigated. These approaches are still subject to elevated levels of genomic off-target mutations outside of the specified gene of interest, which can lead to competitive means of selection escape and reduced host fitness.(*10*–*12*) These limitations can be mitigated by using an orthogonal replication system where a highly error-prone DNA polymerase maintains a dedicated episome.(*13*–*15*) Here, mutation rates are not subject to the limitations imposed by the error-threshold for genome maintenance, and mutagenesis is constricted to a defined replicon thus minimizing selection escape *via* off-target mutations.

The first orthogonal replication system, OrthoRep, was developed by minimizing extranuclear plasmids of yeast that are replicated by an orthogonal cytosolic protein-primed DNA polymerase.(*13*) Subsequent high-throughput engineering of this DNA polymerase yielded clones with 100,000-fold increased mutation rates of 10^−5^ spb *in vivo*.(*12*) The prowess of this highly mutagenic system has been demonstrated by evolving enzymes, antibodies, and entire metabolic pathways.(*16*–*18*) However, in the context of directed evolution, the majority of genetic tools are not directly compatible with yeast, but have been developed for *Escherichia coli* (*E. coli)*, the workhorse organism of synthetic biology.(*19*) More importantly, an orthogonal replication system in *E. coli* will benefit from faster generation times (20–30 min) and higher cell densities (10^9^–10^10^ ml^-1^) compared to yeast (∼1.5–2.5 h; 10^7^–10^8^ cells ml^-1^), thus enabling significantly accelerated evolution experiments at the same mutation rate (∼500-fold more mutations for a given time and culture volume).(*20*) Recently, a bacterial replication system EcORep based on the protein-primed DNA polymerase of the lytic bacteriophage PDR1 has been developed for *E. coli*.(*15*) However, the mutation rates that can stably be maintained in the disclosed system (∼2.0 × 10^−7^ spb) were only modestly enhanced over genomic levels, effectively mutating only 0.02% of a hypothetical 1 kb gene each generation.(*20*) As such, the development of a highly mutagenic orthogonal replication systems with high transformation efficiencies in *E. coli* for continuous hypermutation and accelerated evolution remains an important challenge in the development of continuous evolution systems.

### Establishing an orthogonal replication system based on the bacteriophage T7 replisome

While no natural orthogonal replisome-replicon pairs exist for *E. coli*, there are lytic coliphages that encode for their own replisome.(*21*) Thus, to establish an orthogonal replisome-replicon pair, we capitalized on the vast body of literature detailing the replisome of the lytic bacteriophage T7 (Fig. 1).(*22*–*24*) The replication of the linear double stranded genome of T7 phage is initiated by priming of its replisome by T7 RNA polymerase (gp1) at a T7 origin of replication (Fig. 1A).(*25*) T7 origins of replication rely on the orthogonality of the T7 RNA polymerase/T7 promoter pair to *E. coli* polymerase/promoter pairs for specificity in recruiting the replisome (Fig. 1B).(*26*) The orthogonal T7 transcript primes the T7 DNA polymerase (gp5), which further recruits the T7 helicase (gp4A) and helicase/primase (gp4B) fusion protein to the transcription bubble. The host factor thioredoxin (*trxA*) binds to the T7 DNA polymerase and serves as a β-sliding clamp. Replisome assembly initiates concerted leading and lagging strand synthesis, which further requires the presence of the T7 single-stranded DNA binding protein (gp2.5) for stabilizing the lagging strand replication loop (Fig. 1C).(*25*)

**Fig. 1.**
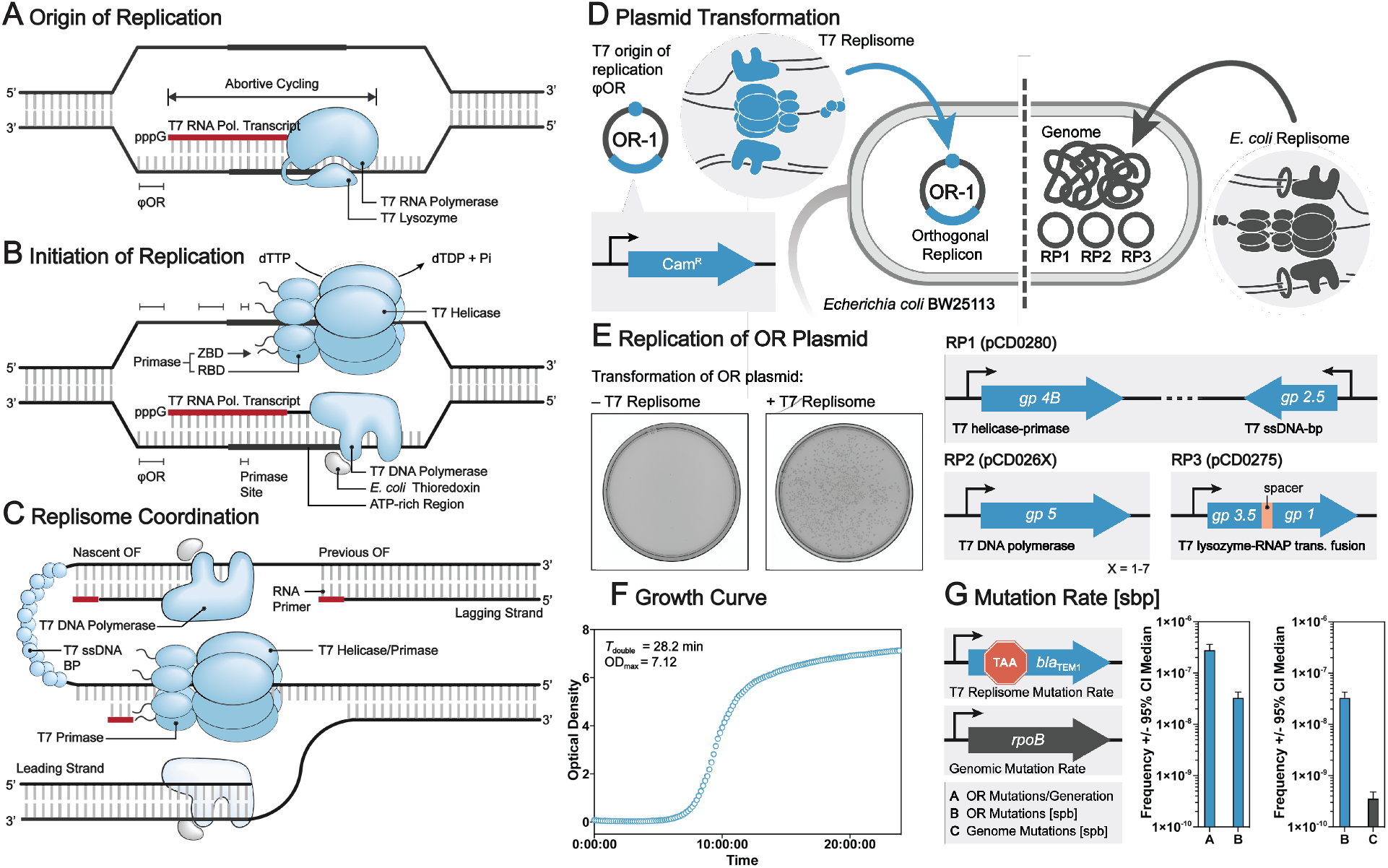
Establishing an orthogonal replication system in *E. coli* based on the bacteriophage T7 replisome. (A) T7 RNA polymerase initiates replication by priming the replisome at the T7 origin of replication. T7 lysozyme stimulates initiation of replication. (B) T7 DNA polymerase is primed by the orthogonal transcript and T7 helicase/primase is recruited to the origin, leading to (C) concerted leading- and lagging-strand synthesis. T7 ssDNA-binding protein stabilizes the replication loop and *E. coli* thioredoxin serves as a sliding clamp to increase T7 DNA polymerase processivity. (D) Transformation of a plasmid with a selection marker and a T7 origin of replication is only replicated by *E. coli* strains that (E) provide the T7 replisome proteins *in trans*. (F) T7 origin plasmids are stably maintained, and transformed strains show robust growth with a doubling time of 28 min and a final OD_600_ of 7.2. (G) The Δ28 mutant of T7 DNA polymerase displays an elevated mutation rate of 3.31 × 10^−8^ spb on the origin replicon (pOR-1), without increasing genomic mutation rate (3.6 ×10^−10^ spb).

To generate an orthogonal T7 replication system in *E. coli*, we generated a strain based on *E. coli* BW25113 (parent strain of Keio collection) in which all replisome genes are supplied *in trans* using plasmid-based expression systems (supplementary material, table S1).(*27*) As described in the literature, we observed severe toxicity for expression of gp4A (helicase), but less toxicity for gp4B (helicase-primase fusion).(*28, 29*) Since gp4A is not essential for replication of T7 phage, we removed the internal start codon of its open reading frame. Expression and functionality of the exogenous proteins was confirmed by infection of the replisome strain with four T7 phages, each with one of the essential replisome genes knocked out. While replication of each knock-out phage proved successful, initial attempts at transforming the replisome strain with a circular plasmid comprising both the primary T7 origin of replication proved unsuccessful.(*30*)

We reasoned that additional T7 genes might be required to establish an orthogonal replication system, but not strictly required for phage propagation.(*31*) While not strictly essential to replication, phages with a T7 lysozyme knockout (gp3.5) are known to display severe replication deficiencies.(*32*) T7 lysozyme is a bifunctional enzyme that displays both cysteine hydrolase activity on *N*-acetylmuramoyl-L-alanine amide bonds of peptidoglycan (major constituent of the bacterial cell wall), as well as inhibition of transcription by T7 RNA polymerase.(*33, 34*) The concerted expression of T7 lysozyme with the other replication proteins (Class II genes of bacteriophage T7, supplementary material fig. S1) implies a regulatory function, in which its accumulation seems to signify a switch from transcription to replication in the phage life cycle.(*35*) Mechanistically, this can be rationalized based on the mechanism of inhibition by T7 lysozyme, which stabilizes T7 RNA polymerase in its binding conformation and prevents the switch to the elongation conformation required for gene transcription.(*36*–*39*) This leads to abortive cycling around the RNA polymerase binding site and to the formation of short RNA ‘primers’ in lieu of the transcription of full genes. The constant opening of the DNA double helix by the transcription bubble enables priming of the T7 DNA polymerase and stimulates assembly of the T7 helicase/primase at the origin.

To maximally benefit from this regulatory effect, we introduced catalytically inactive lysozyme (C131S, hydrolase deficient) as a translational fusion to the T7 RNA polymerase.(*40*) Based on literature precedent, we changed the origin of replication from the primary origin of replication to a secondary origin *φ*OR for which replication stimulation by T7 lysozyme was reported to be highest (Fig. 1A).(*35*) Finally, we chose a previously described exonuclease deficient clone of T7 DNA polymerase (Δ28, T7 Sequenase 2) for initial studies as it had been utilized preferentially in reports on replisome-based *in vitro* replication assays.(*41*–*43*) Implementation of these changes to the replisome strain enabled transformation of a plasmid carrying a T7 origin of replication (pOR-1; supplementary material, table S1), maintaining stable replication for more than 2 weeks (Fig. 1D and E). The strain shows robust growth (*T*_double_ = 29 min) and reaches high cell densities (OD_600_: 7.2; ∼7 × 10^9^ cells ml^-1^) in liquid culture (Fig. 1E). Mutation rates of the initial replisome strain were determined by fluctuation analysis, and showed increased mutagenesis of the T7 origin plasmid (2.81 × 10^−7^ spb per generation) without affecting the genomic mutation rate of the host (3.66 × 10^−10^ spb; supplementary materials, table S4). The copy number of the T7 origin plasmid was determined to be 8.5 based on quantitative PCR, which translates to an adjusted mutation rate of 3.31 × 10^−8^ spb (Fig. 2G, supplementary material, Table S3). Finally, the transformation efficiency of the circular replicon plasmid into the replisome strain was determined to be 2.4 × 10^10^ cfu μg^-1^, orders of magnitude higher than the transformation efficiency for linear plasmids of alternative orthogonal replication systems (i.e., OrthoRep and EcORep).(*12, 15*)

**Fig. 2.**
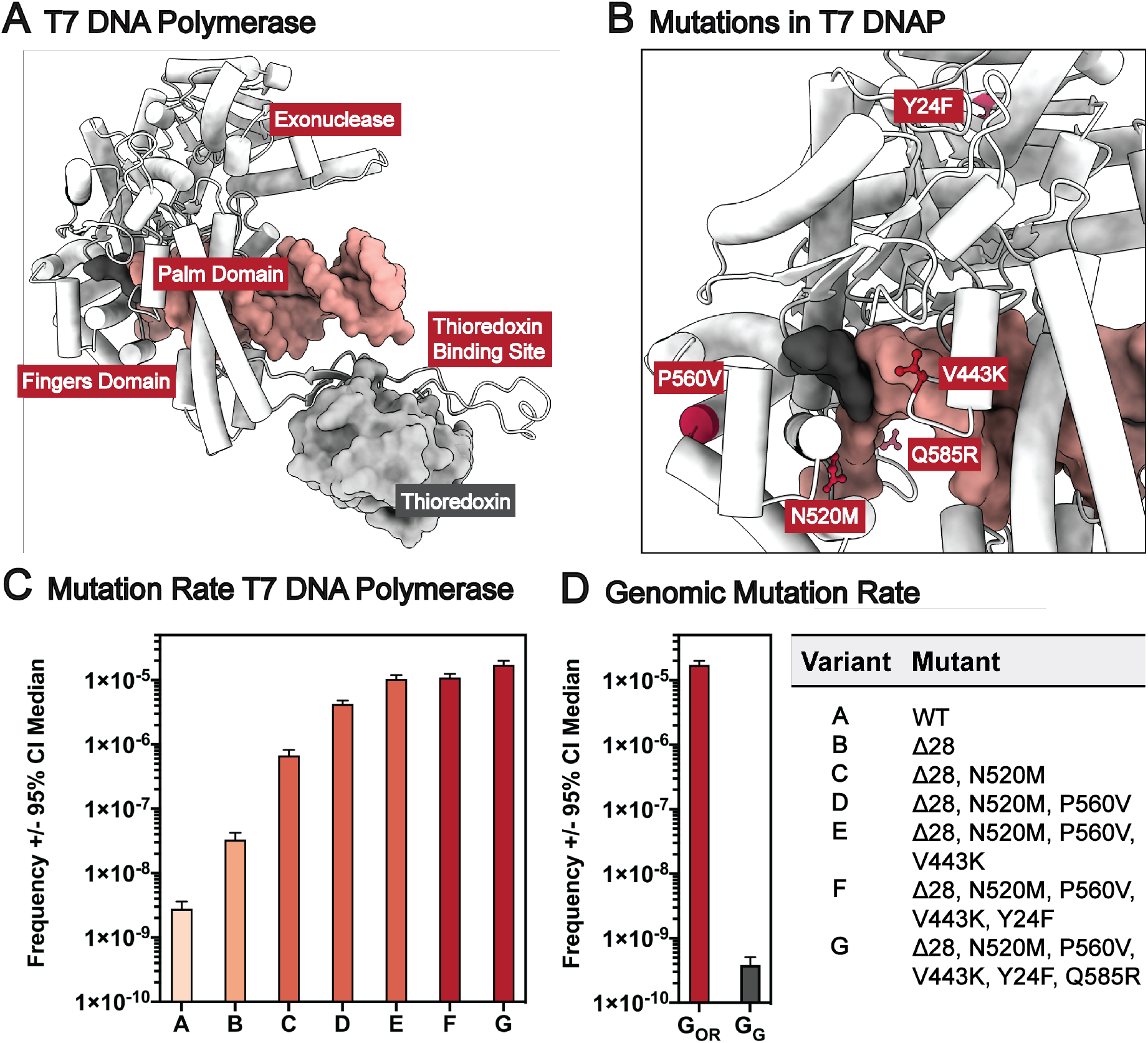
Directed evolution of a mutagenic T7 DNA polymerase. (A) Structure of T7 DNA polymerase (white piped helices) bound to DNA (orange) and its thioredoxin cofactor (light gray). Domains of the polymerase (3’-5’-proofreading exonuclease domain, fingers domain, palm domain, and thioredoxin-binding site) are labeled. (B) Mutations in T7 DNA polymerase that decrease base pairing fidelity (Y24F, V443K, N520M, P560V, Q585R) are highlighted in red (ball and stick representation). (C) Mutation rates of T7 DNA polymerase mutants as determined by fluctuation analysis. Reversion rates are converted to mutation rates by fitting the data to the Luria-Delbruck function using the Ma-Sandri-Sarkar Maximum Likelihood Estimator. (D) Comparison of the mutagenesis by T7 DNA polymerase mutant G on the orthogonal replicon (pOR-3) to the genomic mutation rate shows five orders of magnitude increase in targeted mutagenesis and no off-target activity.

### Engineering T7 DNA Polymerase Mutants with Increased Mutation Rate

The *in vivo* mutation rate of bacteria is controlled by three distinct mechanisms that collectively account for the high fidelity (10^−9^ to 10^−10^ spb) of DNA replication. Specifically, base selection in the T7 DNA polymerase active site accounts for a fidelity of ∼10^−5^ spb, 3’-5’ exonucleolytic proofreading further increases fidelity by ∼10^−2^ spb, and host mismatch repair is responsible for the remaining ∼10^−3^ spb of fidelity. In our efforts to devise a mutagenic orthogonal replisome, we began with a T7 DNA polymerase mutant for which the 3’-5’ proofreading exonuclease activity was removed using a previously described 28 amino acid mutant (Δ28, Fig. 2A). To approximate the extent to which this deletion contributed to mutagenesis *in vivo*, we carried out fluctuation analysis on both the WT polymerase (1.4 × 10^−9^ spb), as well as on the Δ28 mutant (3.31 × 10^−8^ spb), yielding an increase in mutation rate of 25-fold (Fig. 2B). This is in agreement with the magnitude of reported increases in mutation rate *in vitro* (WT: 2.2 × 10^−6^ spb, Δ28 5.4 × 10^−5^ spb, 24.5-fold). Note, the fidelity is ∼3 orders of magnitude higher *in vivo* due to proof-reading by host mismatch repair.(*44*)

Since DNA mismatch repair cannot be altered without compromising host fitness, we next targeted the fidelity of base selection in the polymerase active site to further increase mutagenicity. Initial efforts focused on the identification of sites that contribute to changes in fidelity of homologous polymerases (e.g., *T. aquaticus polA, E. coli polA*, and *S. cerevisiae mip1*), and homology mapping of the corresponding residues onto T7 DNA polymerase. On the basis of previous reports, we targeted 11 sites of T7 DNA polymerase and generated single site-saturation libraries at each position (i.e., R429X, V443X, R444X, L479X, E480X, N520X, A521X, K522X, T523X, F524X, Y530X, P560X, and N611X). We next transformed a replisome strain deficient in T7 DNA polymerase (T7 _Rep-Δgp5_, supplementary material, table S2) with each site-saturation library and carried out reversion assays in which the polymerase libraries were challenged to revert a premature ochre stop codon in the TEM-1 β-lactamase gene (E26*_*ochre*_) on the T7 origin plasmid (pOR-3, supplementary material, table S1), thus restoring antibiotic resistance. After demonstrating survival on the maintenance antibiotic to select for functional DNA polymerases, a reversion selection for mutagenicity on carbenicillin was performed. Surviving clones were sequenced using Illumina amplicon sequencing. Clones that were found to be enriched in the reversion assays were individually subjected to a fluctuation analysis which identified three optimal mutations (N520M, P560V, and V443K, Fig. 2B) that we mapped onto the Δ28 mutant. The mutations at the three identified sites are hypothesized to target distinct mechanisms for increasing the mutation rate. N520M is a mutation in the O-helix, which is in direct contact with the incoming nucleotide in the active site. Mutations at this site likely decrease geometric constraints that contribute to increased base pairing selectivity. P560V is located in a conformationally flexible region of the fingers domain that contacts the O-helix. Mutations in these ‘hinge’ loops that connect the alpha helices of this domain are known to affect the kinetics of conformational changes upon binding of the correct/incorrect nucleotides, thus impacting fidelity. Finally, V443K is a mutation at a site that is in direct contact with the DNA double helix at the replication fork. Such mutations likely decrease base pairing constraints by increasing the flexibility at the replication fork, which minimizes strain due to pairing of mismatched bases. The identified mutations increased the mutation rate of of exonuclease deficient T7 DNAP to 6.75 × 10^−7^ spb for the best single mutant, 4.26 × 10^−6^ spb for the best double mutant, and 1.04 × 10^−5^ spb for the best triple mutant (Fig. 2C).

Finally, we constructed a random mutagenesis library by error-prone PCR on top of the polymerase triple mutant and subjected it to the same antibiotic reversion assay. Sequencing to identify codon positions at which mutations were enriched yielded an additional 7 sites for which individual single-site saturation libraries were constructed and screened (Y24X, P84X, 118X, R171X, 433X, L437X, Q585X, and 678X). The two most beneficial mutations identified were Y24F and Q585R. Notably, the first site is located in the exonuclease domain and was found to reverse growth deficiencies observed for the single-, double- and triple mutants (supplementary material, fig. S5-S10, table S5) but did not show a change in mutation rate (1.1 × 10^−5^ spb). The last mutation was found in the Palm domain of the polymerase and yielded a mutation rate of 1.73 × 10^−5^ spb for the quintuple mutant (Fig. 3C; supplementary material, table S3).

Orthogonality to the genome by fluctuation analysis using rifampicin resistance assays was confirmed for the final mutant, which displayed a genomic mutation rate of 3.9 × 10^−10^ spb (Fig. 2D; supplementary material, table S4). Stability of the quintuple mutant was further confirmed by passaging the strain for two weeks, after which sequencing of the quintuple mutant of the replisome strain showed no changes in sequence.

### Continuous Evolution of TEM-1 β-Lactamase

Next, to demonstrate the utility of the mutagenic T7 replisome system for evolving protein function, we applied it to the continuous evolution of TEM-1 β-lactamase for an expanded substrate scope and improved activity against monobactam (aztreonam) and cephalosporin antibiotics (cefotaxime, ceftazidime, and cefepime, Fig. 3A-C). To this end, the most mutagenic (T7_rep-7_; quintuple mutant) replisome strain was transformed with a T7 origin plasmid encoding chloramphenicol resistance (Cam^R^) as a maintenance antibiotic marker, as well as the *bla*_TEM-1_ β-lactamase gene (Fig. 3A). Initial plating of the strain on antibiotic concentrations that approximate the MIC of TEM-1 against the respective antibiotics of interest (0.1 μg ml^-1^ for aztreonam, cefotaxime, and cefepime; 0.4 μg ml^-1^ for ceftazidime) served as the starting point for the evolution experiment. For each experiment 96 colonies were picked and the cells were passaged under increasing concentrations of antibiotic for 6 days with an overall increase in concentration of ≥1000-fold (100 μg ml^-1^ for aztreonam, cefotaxime, cefepime; 500 μg ml^-1^ for ceftazidime; Fig. 3B). At the end of the experiment, the coding sequences of the β-lactamase genes, as well as the *p*_AmpR_ promoter region were amplified by PCR and analyzed by sequencing (Fig. 3D, supplementary material, fig. S12-S14).

**Fig. 3.**
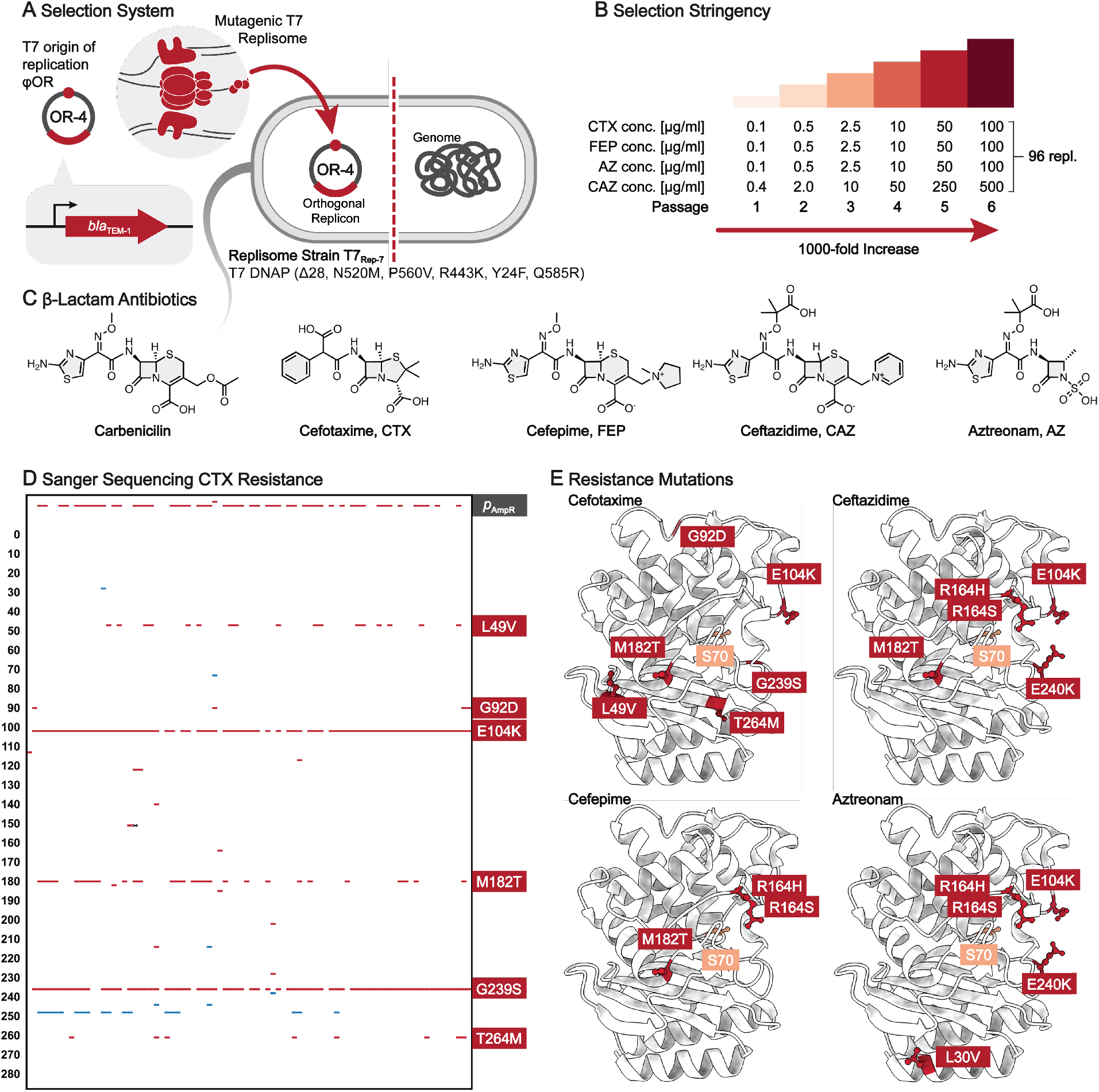
Continuous evolution of TEM-1 β-Lactamase. (A) TEM-1 β-Lactamase is continuously mutagenized by the mutagenic T7 replisome (T7_rep-7_, quintuple mutant). (B) Selection for expanded substrate scopes is achieved by passaging cells (96 replicates) in the presence of increasing amounts of cephalosporin and monobactam antibiotics. (C) Chemical structures of carbenicillin, cephalosporin β-lactams (cefotaxime, cefepime, ceftazidime), and the monobactam aztreonam. (D) Sanger sequencing for 96 cefotaxime resistance evolution experiments after 1000-fold increase in antibiotic concentration (missense mutations are marked in red, silent mutations in blue). (E) Structure of TEM-1 β-lactamase with mutations that multiple replicates converge upon are highlighted in red, the catalytic S70 residue in orange.

Convergence of the mutations across the 96 replicates was observed for all antibiotics, with each of the enriched mutations having previously been described in clinical isolates of extended spectrum β-lactamases and/or in laboratory evolution experiments.(*45, 46*) Both the nature, and the combination of mutations differ for the four distinct β-lactam antibiotics. In addition, the entire 96-well plate for each evolution experiment was pooled after each passage and subjected to nanopore amplicon sequencing to elucidate the order and establish a time-course of when mutations occur (Fig. 4, A and B, and supplementary material fig. S15 and S16). For instance, the evolution of cefotaxime resistance (Fig. 4A) begins to enrich sequences comprising the G238S substitution from the first passage, in agreement with the substitution being associated with increased hydrolysis of cefotaxime. After passage 3, only clones that acquire a secondary mutation, E104K, survive. E104K is generally associated with mutations G238S or R164H/S (see below) where in combination it drastically increases resistance to cefotaxime, or ceftazidime and aztreonam, respectively.(*45, 47*) Starting at passage 4, mutations that increase the strength of the weak promoter *p*_AmpR_ are observed. Finally, over passages 4, 5, and 6 one of four accessory mutations are observed for the majority of cultures (M182T, L49M, T265M, or G92D), all of which have been found as secondary mutations in both clinical isolates and evolution experiments.(*45*) These mutations are more distant from the active site and have no direct effect on catalytic activity. M182T increases thermodynamic stability and helps prevent aggregation. Similarly, G92D and T265M were identified in evolution campaigns that select for stabilizing and compensatory mutations.(*45*) Finally, the role of L49M is unknown, but the fact that it is observed frequently and specifically in the context of cefotaxime resistance supports an adaptive effect.(*45*)

**Fig. 4.**
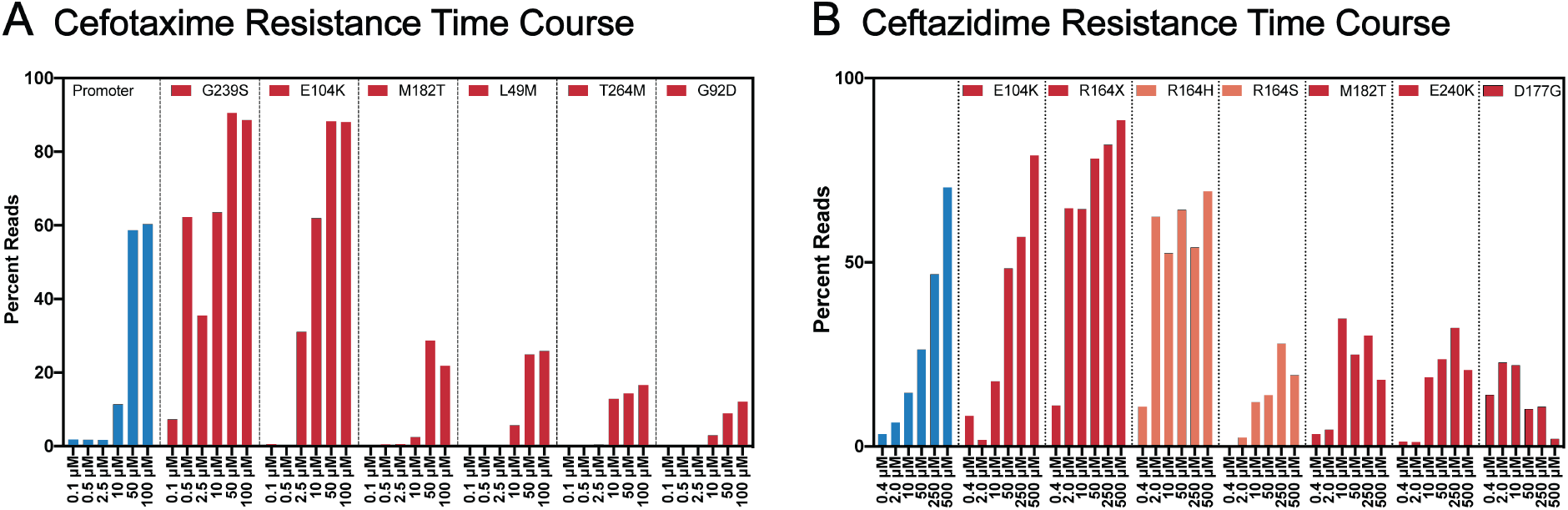
Time course of β-lactamase evolution experiments. The time course for evolution of (A) cefotaxime and (B) ceftazidime resistance was interrogated by deep-sequencing of all 96 replicates after each passage (once per day). Mutations in the promoter are depicted in blue, mutations in the TEM-1 reading frame are depicted in red and orange.

In the ceftazidime resistance time course experiment, mutations in the promoter region are observed from the outset (Fig. 5B). This can be attributed to the higher initial activity against this β-lactam, making increased expression levels a viable solution for improved fitness. Additionally, mutations R164H/S (increases accessibility of β-lactams with larger side-chains to the active site) are enriched from the outset, along with the aforementioned E104K secondary mutation. After passage 3, additional mutations (i.e., M182T, E240K, and D179G) are observed. E240K is a secondary mutation that is commonly observed in combination with R164S/H, where it significantly increases activity against ceftazidime and aztreonam.(*45, 47*) D179G has been observed in evolution campaigns using ceftazidime as a selective agent.(*45*) Over the last three passages, the percentage of D179G mutations is steadily decreasing, indicating that the mutation is being outcompeted at higher antibiotic concentrations. It is interesting to note that this one-week experiment perfectly recapitulates the vast body of work on extended-spectrum β-lactamases collected over decades, highlighting the power of the T7 evolution system in interrogating complex evolutionary fitness landscapes.

## Discussion

We have established an orthogonal replication system in *E. coli* based on the replisome of bacteriophage T7. Directed evolution of the T7 DNA polymerase for increased mutation rates yielded a replisome strain that can be continuously passaged at a mutation rate of 1.7 × 10^−5^ spb – two orders of magnitude higher than that of the best reported orthogonal replication system in *E. coli* to date. The combination of the high mutagenesis rate, the fast growth rate of *E. coli*, as well as the ease with which both the *E. coli* and the circular replicon plasmid can be integrated into standard molecular biology workflows differentiates this system from OrthoRep and EcORep. In comparison to PACE, another highly mutagenic continuous evolution system in *E. coli*, our system benefits from simplicity and the lack of specialized equipment that is required to run evolution campaigns, as well as from the ability to autonomously run continuous evolution campaigns in a large number of replicates.(*48, 49*) Looking forward, the helicase-dependent concerted leading- and lagging-strand synthesis of the T7 replisome, unlike helicase-independent replication modules of previous orthogonal replication systems, enables high transformation efficiencies of circular plasmids (2.4 × 10^10^ cfu μg^-1^) and thus the integration of pre-diversified libraries (e.g., site-saturation libraries) and continuous mutagenesis into a single workflow. This will be particularly beneficial for evolving fundamentally new as opposed to improved activities, as well as for investigating vast fitness landscapes through divergent, as opposed to convergent evolution campaigns.

## Acknowledgments

The authors thank Kristen Williams for assistance with manuscript preparation and submission.

## Author contributions

Conceptualization: CSD, PGS,

Methodology: CSD, PGS

Investigation: CSD, PJS, CR, DAD

Funding acquisition: PGS

Supervision: CSD, PGS

Writing – original draft: CSD, PGS

Writing – review & editing: CSD, PJS, CR, DAD, PGS

## Data and materials availability

The authors have filed patent applications on T7 replication system. Plasmids listed in Table 1 are available from the authors upon reasonable request.

## Supplementary Materials

Materials and Methods

Figs. S1 to S16

Tables S1 to S5

References (*50–54*)

## References

1. J. W. Drake, A constant rate of spontaneous mutation in DNA-based microbes. Proc. Natl. Acad. Sci. 88, 7160–7164 (1991).

2. C. K. Biebricher, M. Eigen, What is a quasispecies? Quasispecies concept Implic. Virol., 1–31 (2006).

3. M. S. Packer, D. R. Liu, Methods for the directed evolution of proteins. Nat. Rev. Genet. 16, 379–394 (2015).

4. F. H. Arnold, Design by directed evolution. Acc. Chem. Res. 31, 125–131 (1998).

5. A. J. Simon, S. d’Oelsnitz, A. D. Ellington, Synthetic evolution. Nat. Biotechnol. 37, 730– 743 (2019).

6. W. P. C. Stemmer, Rapid evolution of a protein in vitro by DNA shuffling. Nature 370, 389–391 (1994).

7. D. M. Hillis, J. J. Bull, M. E. White, M. R. Badgett, I. J. Molineux, Experimental phylogenetics: Generation of a known phylogeny. Science 255, 589–592 (1992).

8. B. C. Dickinson, M. S. Packer, A. H. Badran, D. R. Liu, A system for the continuous directed evolution of proteases rapidly reveals drug-resistance mutations. Nat. Commun. 5 (2014).

9. M. Camps, J. Naukkarinen, B. P. Johnson, L. A. Loeb, Targeted gene evolution in Escherichia coli using a highly error-prone DNA polymerase I. Proc. Natl. Acad. Sci. U. S. A. 100, 9727–9732 (2003).

10. J. J. Bull, R. Sanjuan, C. O. Wilke, Theory of lethal mutagenesis for viruses. J. Virol. 81, 2930–2939 (2007).

11. A. J. Herr, M. Ogawa, N. A. Lawrence, L. N. Williams, J. M. Eggington, M. Singh, R. A. Smith, B. D. Preston, Mutator suppression and escape from replication error–induced extinction in yeast. PLoS Genet. 7, e1002282 (2011).

12. A. Ravikumar, G. A. Arzumanyan, M. K. A. Obadi, A. A. Javanpour, C. C. Liu, Scalable, continuous evolution of genes at mutation rates above genomic error thresholds. Cell 175, 1946–1957 (2018).

13. A. Ravikumar, A. Arrieta, C. C. Liu, An orthogonal DNA replication system in yeast. Nat. Chem. Biol. 10, 175–177 (2014).

14. R. Tian, R. Zhao, H. Guo, K. Yan, C. Wang, C. Lu, X. Lv, J. Li, L. Liu, G. Du, Engineered bacterial orthogonal DNA replication system for continuous evolution. Nat. Chem. Biol. 19, 1504–1512 (2023).

15. R. Tian, F. B. H. Rehm, D. Czernecki, Y. Gu, J. F. Zürcher, K. C. Liu, J. W. Chin, Establishing a synthetic orthogonal replication system enables accelerated evolution in E. coli. Science 383, 421–426 (2024).

16. G. Rix, E. J. Watkins-Dulaney, P. J. Almhjell, C. E. Boville, F. H. Arnold, C. C. Liu, Scalable continuous evolution for the generation of diverse enzyme variants encompassing promiscuous activities. Nat. Commun. 11, 5644 (2020).

17. A. Wellner, C. McMahon, M. S. A. Gilman, J. R. Clements, S. Clark, K. M. Nguyen, M. H. Ho, V. J. Hu, J.-E. Shin, J. Feldman, Rapid generation of potent antibodies by autonomous hypermutation in yeast. Nat. Chem. Biol. 17, 1057–1064 (2021).

18. E. D. Jensen, F. Ambri, M. B. Bendtsen, A. A. Javanpour, C. C. Liu, M. K. Jensen, J. D. Keasling, Integrating continuous hypermutation with high-throughput screening for optimization of cis, cis-muconic acid production in yeast. Microb. Biotechnol. 14, 2617– 2626 (2021).

19. J. E. Cronan, Escherichia coli as an experimental organism. eLS (2014).

20. R. L. Williams, C. C. Liu, Accelerated evolution of chosen genes. Science 383, 372–373 (2024).

21. C. Weigel, H. Seitz, Bacteriophage replication modules. FEMS Microbiol. Rev. 30, 321– 381 (2006).

22. C. C. Richardson, Bacteriophage T7: minimal requirements for the replication of a duplex DNA molecule. Cell 33, 315–317 (1983).

23. A. W. Kulczyk, C. C. Richardson, The replication system of bacteriophage T7. Enzym. 39, 89–136 (2016).

24. Y. Gao, Y. Cui, T. Fox, S. Lin, H. Wang, N. de Val, Z. H. Zhou, W. Yang, Structures and operating principles of the replisome. Science 363, eaav7003 (2019).

25. S. J. Lee, C. C. Richardson, Choreography of bacteriophage T7 DNA replication. Curr. Opin. Chem. Biol. 15, 580–586 (2011).

26. S. Tabor, C. C. Richardson, A bacteriophage T7 RNA polymerase/promoter system for controlled exclusive expression of specific genes. Proc. Natl. Acad. Sci. 82, 1074–1078 (1985).

27. T. Baba, T. Ara, M. Hasegawa, Y. Takai, Y. Okumura, M. Baba, K. A. Datsenko, M. Tomita, B. L. Wanner, H. Mori, Construction of Escherichia coli K-12 in-frame, single-gene knockout mutants: The Keio collection. Mol. Syst. Biol. 2, 8–2006 (2006).

28. H. Zhang, S.-J. Lee, A. W. Kulczyk, B. Zhu, C. C. Richardson, Heterohexamer of 56-and 63-kDa gene 4 helicase-primase of bacteriophage T7 in DNA replication. J. Biol. Chem. 287, 34273–34287 (2012).

29. W. Ma, A. Phan, R. Walsh, K. Ye, Building an orthogonal replication system for performing directed evolution in Escherichia coli: A strategic review and a summary of the initial steps in cloning bacteriophage T7 gp4 primase/helicase. JEMI, 1–8 (2015).

30. K. Becker, A. Meyer, T. M. Roberts, S. Panke, Plasmid replication based on the T7 origin of replication requires a T7 RNAP variant and inactivation of ribonuclease H. Nucleic Acids Res. 49, 8189–8198 (2021).

31. H. Saito, S. Tabor, F. Tamanoi, C. C. Richardson, Nucleotide sequence of the primary origin of bacteriophage T7 DNA replication: Relationship to adjacent genes and regulatory elements. Proc. Natl. Acad. Sci. U. S. A. 77, 3917–3921 (1980).

32. W. T. McAllister, H.-L. Wu, Regulation of transcription of the late genes of bacteriophage T7. Proc. Natl. Acad. Sci. 75, 804–808 (1978).

33. B. A. Moffatt, F. W. Studier, T7 lysozyme inhibits transcription by T7 RNA polymerase. Cell 49, 221–227 (1987).

34. X. Cheng, X. Zhang, J. W. Pflugrath, F. W. Studier, The structure of bacteriophage T7 lysozyme, a zinc amidase and an inhibitor of T7 RNA polymerase. Proc. Natl. Acad. Sci. 91, 4034–4038 (1994).

35. X. Zhang, F. W. Studier, Multiple roles of T7 RNA polymerase and T7 lysozyme during bacteriophage T7 infection. J. Mol. Biol. 340, 707–730 (2004).

36. X. Zhang, F. W. Studier, Mechanism of inhibition of bacteriophage T7 RNA polymerase by T7 lysozyme. J. Mol. Biol. 269, 10–27 (1997).

37. D. Jeruzalmi, T. A. Steitz, Structure of T7 RNA polymerase complexed to the transcriptional inhibitor T7 lysozyme. EMBO J. 17, 4101–4113 (1998).

38. A. Kumar, S. S. Patel, Inhibition of T7 RNA polymerase: Transcription initiation and transition from initiation to elongation are inhibited by T7 lysozyme via a ternary complex with RNA polymerase and promoter DNA. Biochemistry 36, 13954–13962 (1997).

39. N. M. Stano, S. S. Patel, T7 lysozyme represses T7 RNA polymerase transcription by destabilizing the open complex during initiation. J. Biol. Chem. 279, 16136–16143 (2004).

40. B. C. Dickinson, M. S. Packer, A. H. Badran, D. R. Liu, A system for the continuous directed evolution of proteases rapidly reveals drug-resistance mutations. Nat. Commun. 5, 5352 (2014).

41. S. Tabor, C. C. Richardson, DNA sequence analysis with a modified bacteriophage T7 DNA polymerase. Proc. Natl. Acad. Sci. 84, 4767–4771 (1987).

42. S. Doublié, T. Ellenberger, The mechanism of action of T7 DNA polymerase. Curr. Opin. Struct. Biol. 8, 704–712 (1998).

43. B. Zhu, Bacteriophage T7 DNA polymerase–sequenase. Front. Microbiol. 5, 89476 (2014).

44. T. A. Kunkel, S. S. Patel, K. A. Johnson, Error-prone replication of repeated DNA sequences by T7 DNA polymerase in the absence of its processivity subunit. Proc. Natl. Acad. Sci. 91, 6830–6834 (1994).

45. M. L. M. Salverda, J. A. G. M. De Visser, M. Barlow, Natural evolution of TEM-1 β-lactamase: Experimental reconstruction and clinical relevance. FEMS Microbiol. Rev. 34, 1015–1036 (2010).

46. R. Fernandes, P. Amador, C. Oliveira, C. Prudêncio, Molecular characterization of ESBL-producing Enterobacteriaceae in northern Portugal. Sci. World J. 2014 (2014).

47. T. Palzkill, Structural and mechanistic basis for extended-spectrum drug-resistance mutations in altering the specificity of TEM, CTX-M, and KPC β-lactamases. Front. Mol. Biosci. 5, 16 (2018).

48. S. M. Miller, T. Wang, D. R. Liu, Phage-assisted continuous and non-continuous evolution. Nat. Protoc. 15, 4101–4127 (2020).

49. K. M. Esvelt, J. C. Carlson, D. R. Liu, A system for the continuous directed evolution of biomolecules. Nature 472, 499–503 (2011).

