## Supplementary Materials for "An Orthogonal T7 Replisome for Continuous Hypermutation and Accelerated Evolution in *E. coli*"

**This PDF file includes:**

Materials and Methods
Figs. S1 to S16
Table S1 to S5
References (*50–54*)

### Materials and Methods

**Culture conditions and reagents.** All *E. coli* cultures were grown in 2YT media unless otherwise specified. Cloning was carried out in *E. coli* DH5a. All replisome strains were constructed in *E. coli* BW25113 (*$F^-$ $LAM^-$*::rrnB3, Δ*lacZ4787, hsdR514,* Δ*(araBAD)567,* Δ*(rhaBAD)568 rph-1*), the Keio collection parent strain.(*50*) Antibiotics were used at the following concentrations unless specified otherwise: Kanamycin (50 µg/ml), stretopmycin (100 µg/ml), gentamycin (20 µg/ml), carbenicillin (100 µg/ml), chloramphenicol (25 µg/ml), and rifampicin (50 µg/ml).

**DNA cloning.** Plasmids used in this study are listed in Table S1. *E. coli* strain NEB5a (NEB) was used for all DNA cloning steps. All primers used in this study were purchased from Integrated DNA Technologies, USA. All enzymes for PCR and cloning were obtained from NEB. Plasmids were assembled by Gibson assembly. To clone the T7 phage replisome constructs (Table S1), DNA fragments encoding the open-reading frames of gp1, gp2.5, gp4, gp3.5, and gp5 were purchased from IDT.

**Growth curves.** Growth curves of the replisome strains (Table S2) transformed with pOR-3 (Table S1) were measured in a 96-well plate format using the BioTek LogPhase 600 microbiology reader. Cultures of 150 µl in volume were grown at 37 °C at a shaking speed of 800 (a.u.). Terrific broth was used as the culture media and was supplemented with antibiotics kanamycin (22.5 µg/ml), stretopmycin (90 µg/ml), gentamycin (18 µg/ml), and chloramphenicol (25 µg/ml). Optical density was recorded every 10 minutes and samples were measured in 12 replicates for each experiment.

**Transformation efficiency determination.** The transformation efficiency is defined as the theoretical number of colony-forming units (cfu's) produced by transforming 1 µg of plasmid DNA into a given volume of competent cells. In practice, this efficiency was calculated by transforming 100 pg of purified plasmid under idealized conditions. Electrocompetent cells of strain $T7_{rep-2}$ were generated by back-diluting an overnight culture 1000-fold, followed by growing the culture (500 ml) to an optical density ($OD_{600}$) of 0.4 at 37°C. 4 cycles of centrifugation (7 min, 3000×g, 16°C) and media exchange with a full culture volume of 10% glycerol in water were used to remove remaining salt from the media. The final $OD_{600}$ of the competent cells was adjusted to ~200. 50 µl of competent cells were mixed with 100 pg of purified DNA of pOR-2 (Table S1) and electroporated using pre-chilled electroporation cuvettes. Cultures were recovered in 250µl of SOC media for 1 hour, and diluted 10-fold before plating 30 µl on pre-warmed plates (37°C), supplemented with kanamycin (45 µg/ml), stretopmycin (90 µg/ml), gentamycin (18 µg/ml), and carbenicilin (100 µg/ml). The effective transformation efficiency was calculated by normalizing the number of cfu's to a hypothetical DNA concentration of 1 µg.

**Site-saturation mutagenesis.** NNK site-saturation libraries were generated by amplifying target sequences using primers for which one codon is randomized (NNK codon), followed by plasmid ligation using Gibson assembly. The cloned libraries were transformed into NEB® 10-beta electrocompetent *E. coli* cells and plasmid DNA was isolated and purified. Codons that were randomized in the coding sequence of T7 DNA polymerase comprise R429, V443, R444, L479, E480, N520, A521, K522, T523, F524, Y530, P560, N611 in plasmid pCD0261 (Table S1; Δ28 exonuclease deficient mutant), and Y24, P84, M146X, R171, A433, L437, Q585, and R678 in the coding sequence of T7 DNA polymerase of plasmid pCD0264 (Table S1; triple mutant). The numbering of all residues is based off of the wild type T7 DNA polymerase sequence.

**Reversion assays of single site-saturation library of T7 DNAP.** Purified DNA of the NNK single site-saturation T7 DNA polymerase library plasmid (see above) was co-transformed with plasmid pOR-3 (Table S1) by electroporation into strain $T7_{Rep\text{-}\Delta gp5}$ (Table S2) and plated on LB agar supplemented with maintenance antibiotics (kanamycin (45 µg/ml), stretopmycin (90 µg/ml), gentamycin (18 µg/ml), and chloramphenicol (25 µg/mL). Cells were scraped and resuspended using Dulbecco's Phosphate Buffered Saline (DPBS) and 1ml to 10 ml of $OD_{600}$ = 1 re-plated onto LB agar plates of appropriate size supplemented with maintenance antibiotics, as well as selection antibiotic (carbenicillin, 100 µg/ml). Survival of clones is contingent upon reversion of a premature ochre stop codon in the TEM-1 β-Lactamase gene on pOR-3 (i.e., statistical selection for mutagenic T7 DNA polymerase clones).

**Illumina amplicon sequencing.** Cells of carbenicilin resistant clones from the single site-saturation libraries (see above; plated on selective media) were scraped using Dulbecco's Phosphate Buffered Saline (DPBS) buffer and their plasmid DNA was purified using the ZymoPURE II Plasmid Midiprep kit. A 500 base pair sequence comprising the single-site NNK library in the T7 DNA polymerase gene was amplified from the isolated DNA (1 µg of template for 50 µl PCR reaction) using primers functionalized with adaptors for Illumina sequencing. The product was PCR purified using the Monarch® PCR & DNA Cleanup Kit (NEB), and the DNA quantified using the qubit DNA fluorometry method. Samples were analyzed using Azenta Genewiz Amplicon EZ sequencing (50,000 reads) and the codon distribution analyzed for enrichment to identify mutagenic mutants.

**Error-prone PCR library reversion assay.** Purified DNA of the error-prone library plasmid (see above) was co-transformed with plasmid pOR-3 (Table S1) by electroporation into strain $T7_{Rep\text{-}\Delta gp5}$ (Table S2) and plated on LB agar supplemented with maintenance antibiotics (kanamycin (45 µg/ml), stretopmycin (90 µg/ml), gentamycin (18 µg/ml), and chloramphenicol (25 µg/mL). Cells were scraped and resuspended using Dulbecco's Phosphate Buffered Saline (DPBS) and 1ml to 10 ml of $OD_{600}$ 1 re-plated onto LB agar plates of appropriate size supplemented with maintenance antibiotics kanamycin, as well as selection antibiotic (carbenicillin, 100 µg/ml). Survival of clones is contingent upon reversion of a premature ochre stop codon in the TEM-1 β-Lactamase gene on pOR-3 (*i.e.*, statistical selection for mutagenic T7 DNA polymerase clones).

**Small-scale pOR-3 fluctuation analysis.** Strain $T7_{Rep\text{-}\Delta gp5}$ was co-transformed with pOR-3 and promising T7 DNAP clones in either pCD0261 (identified from single site-saturation library reversion assays) or pCD0264 (identified from error-prone PCR library reversion assay). The transformation was plated on LB agar supplemented with maintenance antibiotics kanamycin (45 µg/ml), stretopmycin (90 µg/ml), gentamycin (18 µg/ml), and chloramphenicol (25 µg/ml). Single colonies were picked to grow out 8 replicates overnight in a 96-well plate (150µl volume, 37 °C, shaking speed of 800 a.u.). The cultures were diluted $10^5$-fold and grown until they reach stationary phase. Volumes of 0.5 – 100µl were plated onto selective LB agar media supplemented with maintenance antibiotics and selection antibiotic carbenicillin (100 µg/ml) and the number of revertants (cfu's) was quantified. Dilution plates were used to determine the absolute number of cells plated (no selection antibiotic) to calculate mutation rates. Luria-Delbrück fluctuation analysis was carried out by fitting the obtained data using the MSS maximum likelihood method in the Fluctuation AnaLysis CalculatOR (FalCOR) online tool.(*51*)

**Large-scale pOR-3 fluctuation analysis.** Strains $T7_{Rep\text{-}1}$, $T7_{Rep\text{-}2}$, $T7_{Rep\text{-}3}$, $T7_{Rep\text{-}4}$, $T7_{Rep\text{-}5}$, and $T7_{Rep\text{-}6}$, were transformed with pOR-3 and plated on LB agar media supplemented with maintenance antibiotics kanamycin (45 µg/ml), stretopmycin (90 µg/ml), gentamycin (18 µg/ml), and chloramphenicol (25 µg/ml). Single colonies were picked to grow out 24 replicates overnight in a 96-well plate (150µl volume, 37 °C, shaking speed of 800 [a.u.]). The cultures were back-diluted $10^5$-fold and grown until stationary phase. Volumes of 0.5 – 100µl (depending on the mutation rate) were plated onto selective LB agar media supplemented maintenance antibiotics and selection antibiotic carbenicillin (100 µg/ml) and the number of revertants (cfu's) counted. Dilution plates were used to determine the absolute number of cells plated (no selection antibiotic) to calculate mutation rates. Luria-Delbrück fluctuation analysis was carried out by fitting the data using the MSS maximum likelihood method in the Fluctuation AnaLysis CalculatOR (FalCOR) tool.(*2*) To calculate the mutation rate per base pair (spb), the mutation rate was normalized by the copy number of the plasmid (see below) and the number of mutations that yield functional TEM-1 derivatives (7/9 possible mutations of the ochre codon; divide observed mutation rate by 7/3).

**Genomic fluctuation analysis.** $T7_{Rep-2}$ and $T7_{Rep-6}$ were grown out from glycerol stocks overnight in a 96-well plate (45 µg/ml kanamycin, 90 µg/ml stretopmycin, 18 µg/ml gentamycin). The culture was back-diluted $10^4$-fold and 12 replicates grown out to stationary phase in a 96-well plate format (150µl volume, 37 °C, shaking speed of 800 [a.u.]. 100 µl of culture were plated onto selective LB agar media supplemented with maintenance antibiotics kanamycin (45 µg/ml), stretopmycin (90 µg/ml), gentamycin (18 µg/ml), and selection antibiotic rifampicin (50 µg/ml) and the number of revertants (cfu's) counted. Dilution plates were used to determine the absolute number of cells plated (no selection antibiotic) to calculate mutation rates. Luria-Delbrück fluctuation analysis was carried out by fitting the data using the MSS maximum likelihood method in the Fluctuation AnaLysis CalculatOR (FalCOR) tool.(*51*) To calculate the mutation rate per base pair (s.p.b.), the mutation rate was normalized by the number of mutations in the *rpoB* gene that that impart rifampicin resistance (77 known point mutations, divide observed mutation rate by 77/3).(*52*)

**pOR-3 copy number determination.** The copy number of pOR-3 in the different replisome strains ($T7_{Rep-1}$ – $T7_{Rep-7}$) was evaluated by qPCR using the Power SYBR Green PCR Master Mix (Thermo Fisher, Massachusetts, United States) on the QuantStudio 7 Flex Real-Time PCR System, 384-well (Thermo Fisher, Massachusetts, United States). Calibrator plasmid pSR-1 was constructed by cloning *E. coli* gene *dxs* into pUC19 using Gibson assembly to enable absolute quantitation. *dxs* and $bla_{TEM-1}$ were amplified from 10-fold serial dilutions of pSR-1 and a calibration curve was constructed (Figure S11). For each replisome strain ($T7_{Rep-1}$ – $T7_{Rep-7}$), 100 µL of culture was grown to mid log phase in 2YT media supplemented with kanamycin (45 µg/ml), stretopmycin (90 µg/ml), gentamycin (18 µg/ml), and chloramphenicol (25 µg/ml). The cultures were boiled for ten minutes and spun down for 10 min (21.000 × g). The lysate was serial-diluted in 10-fold increments, which were used as template for qPCR. Amplification of $bla_{TEM-1}$ of pOR-3 and *dxs* of the chromosomal DNA was used to determine the absolute copy number of T7 origin plasmid in the replisome strains by the ΔΔCT method, with *dxs* used to normalize the plasmid DNA amount to the genomic copy number of 1.(*53*)

**β-Lactamase nomenclature.** The numbering of residues in TEM-1 β-lactamase was done according to a standardized nomenclature for class A β-lactamases.(*54*)

**β-Lactamase evolution experiments.** Strain $T7_{Rep-7}$ transformed with pOR-4 was grown out overnight in 2YT media supplemented with maintenance antibiotics (45 µg/ml kanamycin, 90 µg/ml stretopmycin, 18 µg/ml gentamycin, 25 µg/ml chloramphenicol. 100 µl of liquid culture was plated on LB agar plates supplemented with both maintenance antibiotics and selection antibiotic (cefotaxime, ceftatzidime, cefepime, and aztreonam) at a concentration approximating the respective minimum inhibitory concentration of the four β-lactams (0.1 µg/ml, 0.4 µg/ml, 0.1 µg/ml, and 0.1 µg/ml, respectively). 96 colonies for each selection were picked and grown out in 96 deep-well plates (1.2 ml culture volume). The concentration of the selection antibiotic was continuously increased 1000-fold in 5 increments. Cultures were back-diluted up to two times per day (once stationary phase is reached, 100-fold dilutions) and the antibiotic concentration increased once per day (for cefotaxime/cefepime/aztreonam: 0.1 µg/ml, 0.5 µg/ml, 2.5 µg/ml, 10 µg/ml, 50 µg/ml, 100 µg/ml; for ceftazidime: 0.4 µg/ml, 2.0 µg/ml, 10 µg/ml, 50 µg/ml, 250 µg/ml, 500 µg/ml).

**β-Lactamase evolution Sanger sequencing.** At the end of each β-Lactamase evolution experiment the entire 96-well plate was spun down, each individual culture was resuspended in 1 ml of molecular biology grade water, and the entire TEM-1 β-Lactamase coding sequence, as well as the $p_{\text{AmpR}}$ promoter were amplified by PCR. The 96 samples were analyzed by Sanger sequencing of the unpurified PCR product.

**β-Lactamase evolution nanopore amplicon sequencing.** After each day of the β-Lactamase evolution experiment, 100µl of each culture of the 96-well plate were combined, 5ml of the mixed culture was spun down, the plasmid DNA was isolated, and the TEM-1 β-Lactamase gene was amplified by PCR using 1 ng of isolated DNA as template. The samples were analyzed by Primordium nanopore amplicon sequencing.

### Supplementary Figures

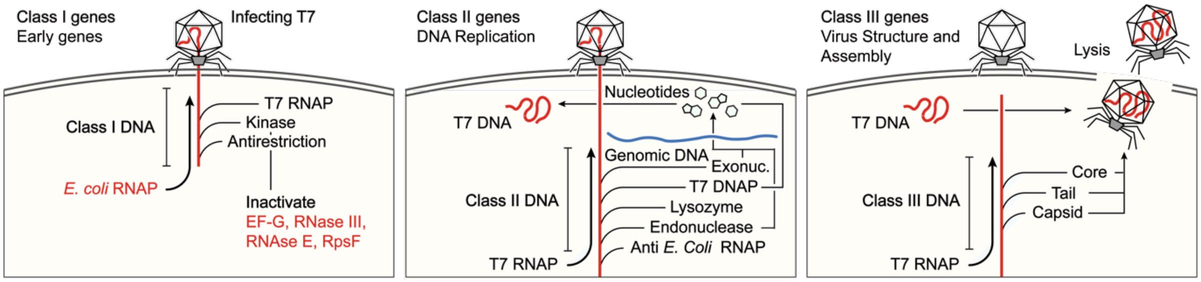


**Figure S1. Bacteriophage T7 life-cycle.** Class I genes encode anti-host function, as well as the T7 RNA polymerase (gp1); Class II genes comprise the T7 genes responsible for replication: ssDNA-binding protein (gp2.5) DNA Polymerase (gp3), DNA helicase and DNA primase (gp4), Class III genes comprise capsid proteins.

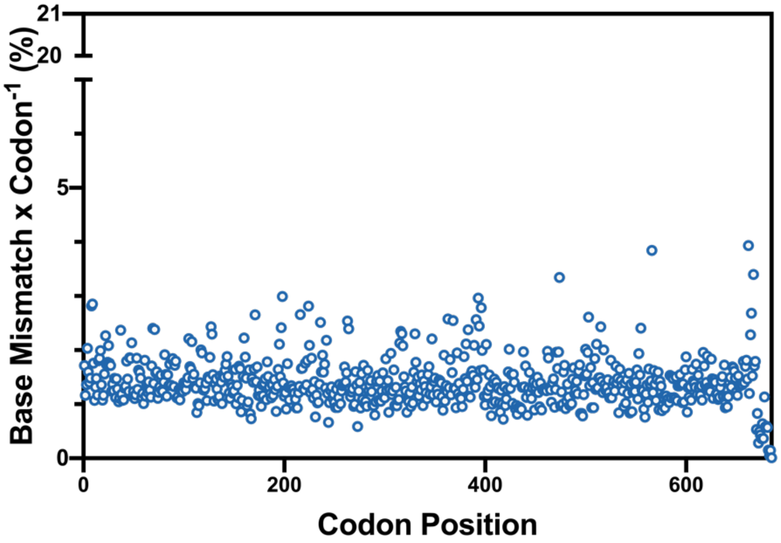


**Figure. S2. Base mismatch per codon of in cloned T7 DNAP error-prone library.** Nanopore sequencing of T7 DNA polymerase library on top of triple mutant (Δ28, N520M, P560V, V443K) in pCD0264. DNA of libraries was purified after transformation of DH5a with the library and plating on solid media (selection for Gentamycin resistance). The coding sequence of T7 DNA polymerase was amplified by PCR, and the product deep-sequenced (Primordium premium PCR, 6000 reads).

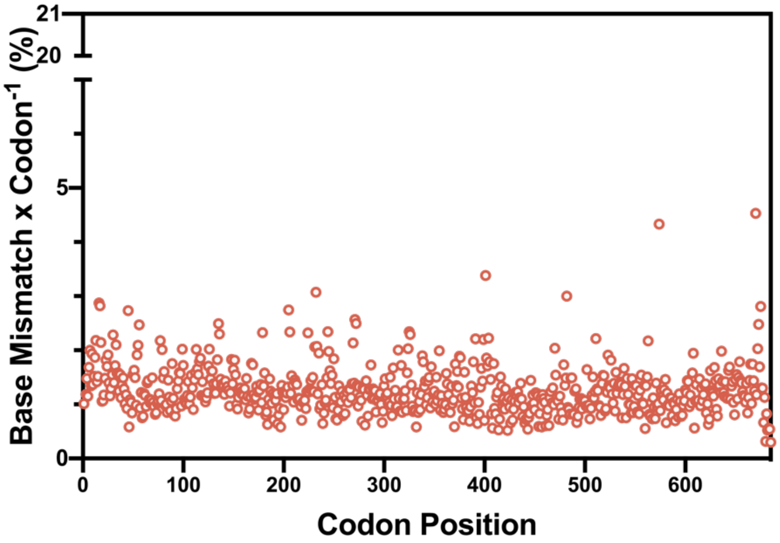


**Figure S3 Base mismatch per codon of survival selection of cloned error-prone T7 DNAP library.** Nanopore sequencing of T7 DNA polymerase library on top of triple mutant (Δ28, N520M, P560V, V443K) in pCD0264 after survival selection for functional DNA polymerase clones: Double transformation of pCD0264 library and pOR-3, followed by selection for chloramphenicol resistance. DNA of libraries was purified afttter selection on solid media, the coding sequence of T7 DNA polymerase amplified by PCR, and the product deep-sequenced (Primordium premium PCR, 6000 reads).

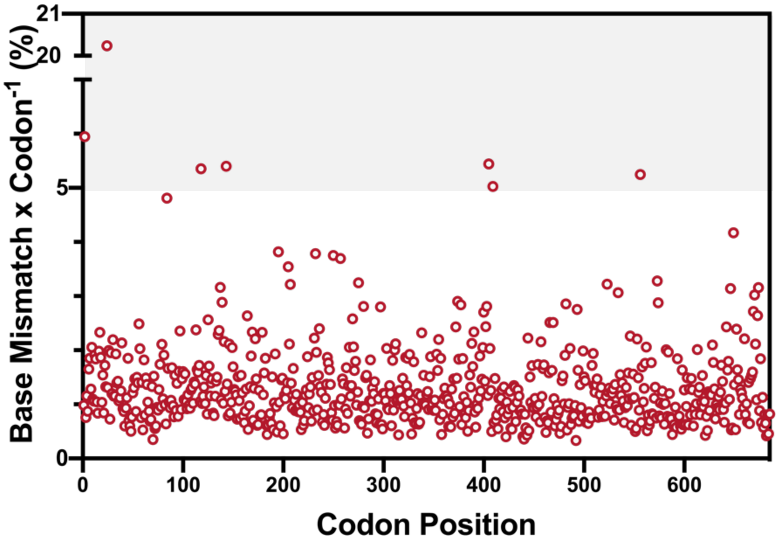


**Figure S4 Base mismatch per codon of mutagenesis selection of cloned error-prone T7 DNAP library.** Nanopore sequencing of T7 DNA polymerase library on top of triple mutant (Δ28, N520M, P560V, V443K) in pCD0264 after selection for mutagenic DNA polymerase clones: Double transformation of pCD0264 library and pOR-3, selection for chloramphenicol resistance). DNA of libraries was purified after selection on solid media, the coding sequence of T7 DNA polymerase amplified by PCR, and the PCR products deep-sequenced (Primordium premium PCR, 6000 reads). Cones that were found to be enriched >5% were chosen (grey background).

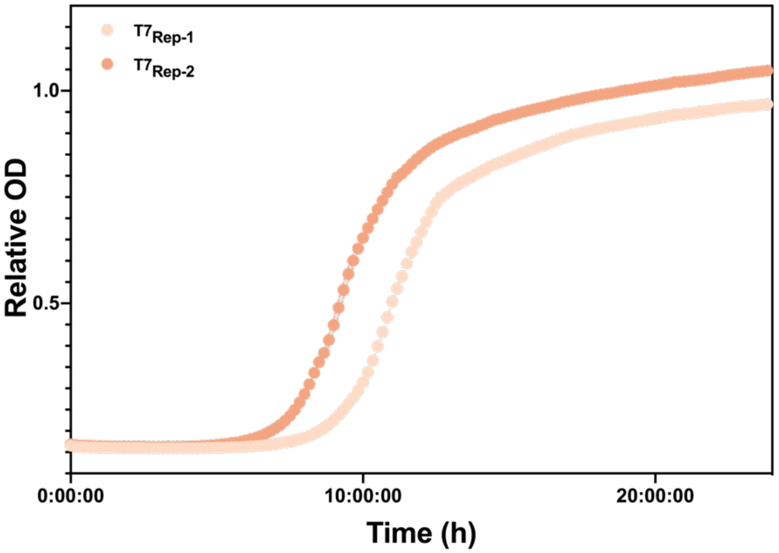


**Figure S5. Growth curve comparison $T7_{Rep-1}$ and $T7_{Rep-2}$.** The strain grows substantially slower when the replisome with the WT DNA polymerase ($T7_{Rep-1}$) maintains the pOR-1 plasmid than when it is maintained by the Δ28 mutant ($T7_{Rep-2}$).

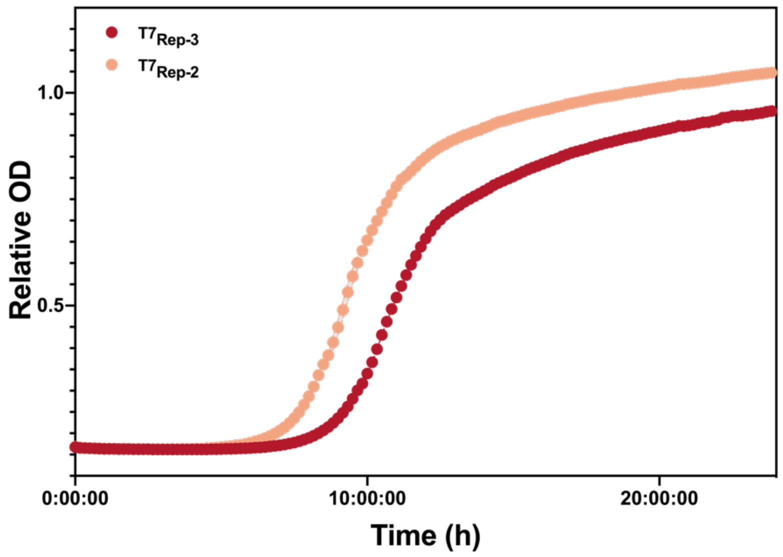


**Figure S6. Growth curve comparison $T7_{Rep-2}$ and $T7_{Rep-3}$.** The strain grows equally fast when the replisome with the single mutant (Δ28, N520M; $T7_{Rep-3}$) DNA polymerase maintains the pOR-1 ($T7_{Rep-3}$) plasmid than when it is maintained by the Δ28 mutant ($T7_{Rep-2}$).

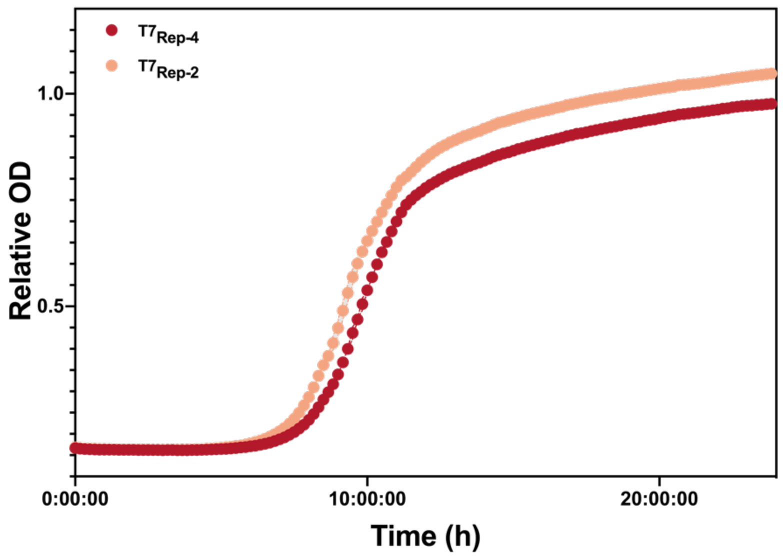


**Figure S7. Growth curve comparison $T7_{Rep-2}$ and $T7_{Rep-4}$.** The strain grows equally fast when the replisome with the double mutant (Δ28, N520M, P560V; $T7_{Rep-4}$) DNA polymerase maintains the pOR-1 plasmid than when it is maintained by the Δ28 mutant ($T7_{Rep-2}$).

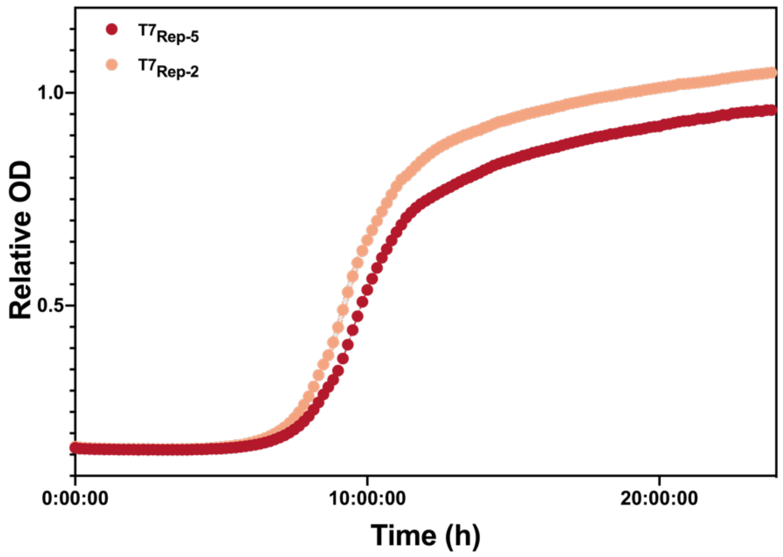


**Figure S8. Growth curve comparison T7$_{Rep-2}$ and T7$_{Rep-5}$.** The strain grows equally fast when the replisome with the triple mutant (Δ28, N520M, P560V; T7$_{Rep-5}$) DNA polymerase maintains the pOR-1 plasmid than when it is maintained by the Δ28 mutant (T7$_{Rep-2}$).

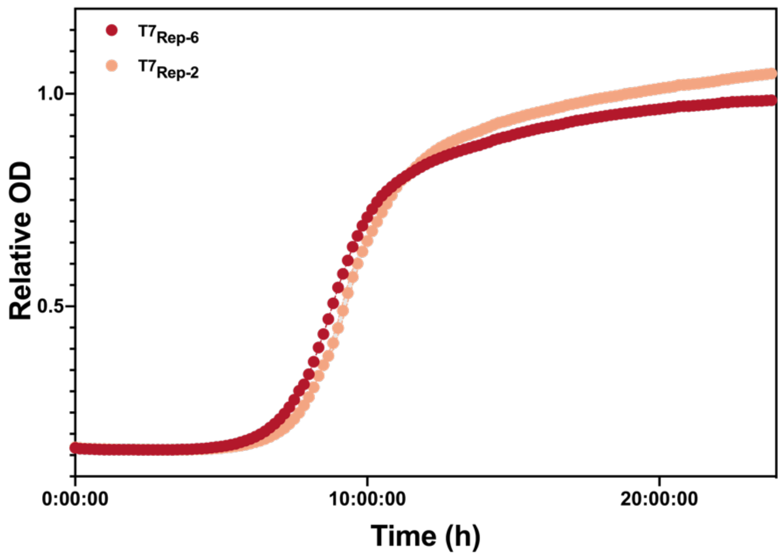


**Figure S9. Growth curve comparison T7$_{Rep-2}$ and T7$_{Rep-6}$.** The strain grows equally fast when the replisome with the quadruple mutant (Δ28, N520M, P560V, Y24F; T7$_{Rep-6}$) DNA polymerase maintains the pOR-1 plasmid than when it is maintained by the Δ28 mutant (T7$_{Rep-2}$).

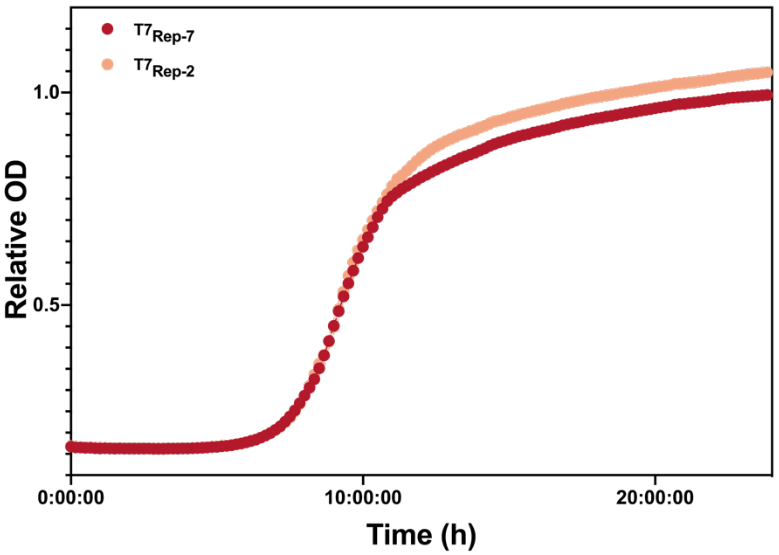


**Figure S10. Growth curve comparison T7$_{Rep-2}$ and T7$_{Rep-7}$.** The strain grows equally fast when the replisome with the quintuple mutant (Δ28, N520M, P560V, Y24F, Q585R; T7$_{Rep-6}$) T7 DNA polymerase maintains the pOR-1 plasmid than when it is maintained by the Δ28 mutant (T7$_{Rep-2}$).

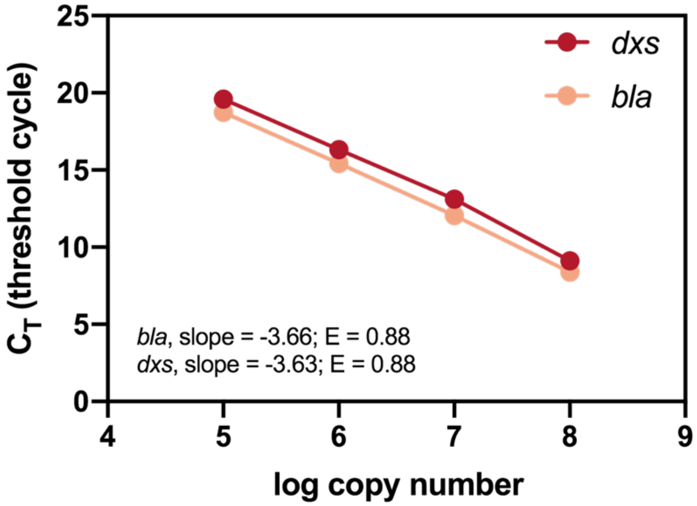


**Figure S11. Calibration curve of amplification efficiency for dxs and TEM-1 genes.** Serial dilutions of pSR1 were amplified for *bla*$_{TEM-1}$ and *dxs* using qPCR to construct a standard curve. Primer efficiency (E) and the difference in threshold cycle (ΔCT) between the genes were determined from the curve and were used to calculate plasmid copy number of the samples.

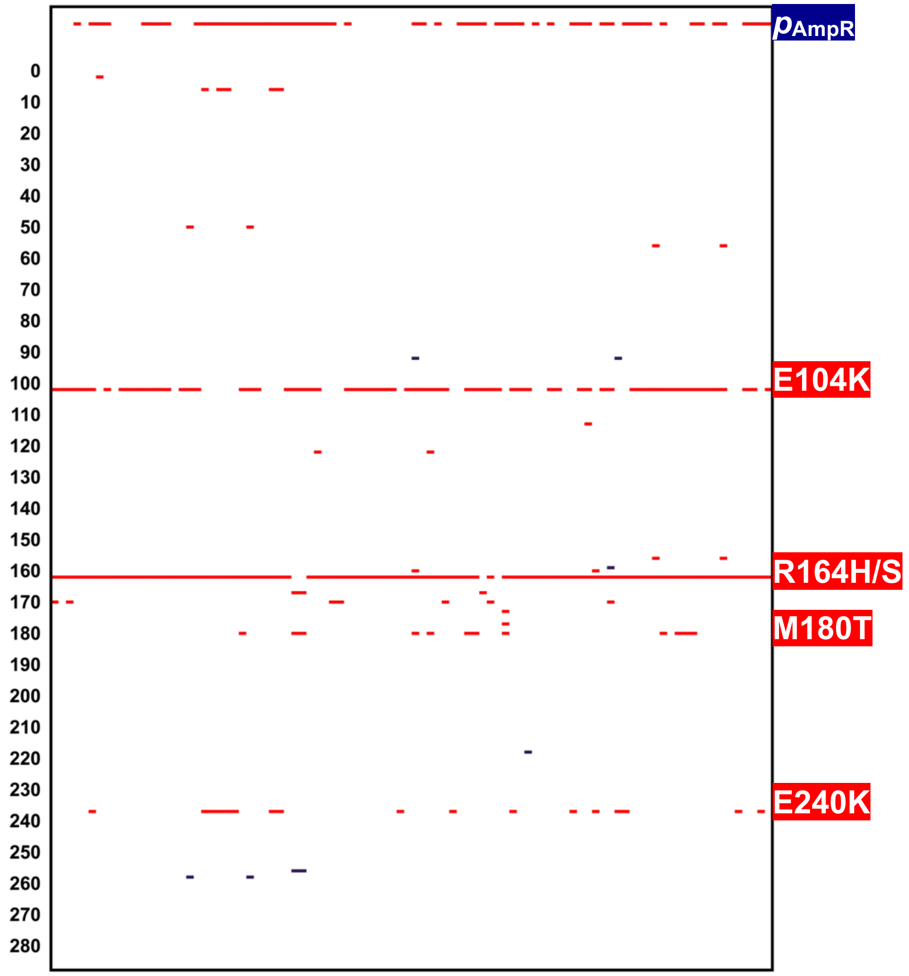


**Figure S12. Sanger sequencing of ceftazidime resistance continuous evolution experiment.** Convergence of sequences in the promoter region ($p_{\text{AmpR}}$), as well as in the coding sequence (E104K, R164H, R164S, M180T, and E240K).

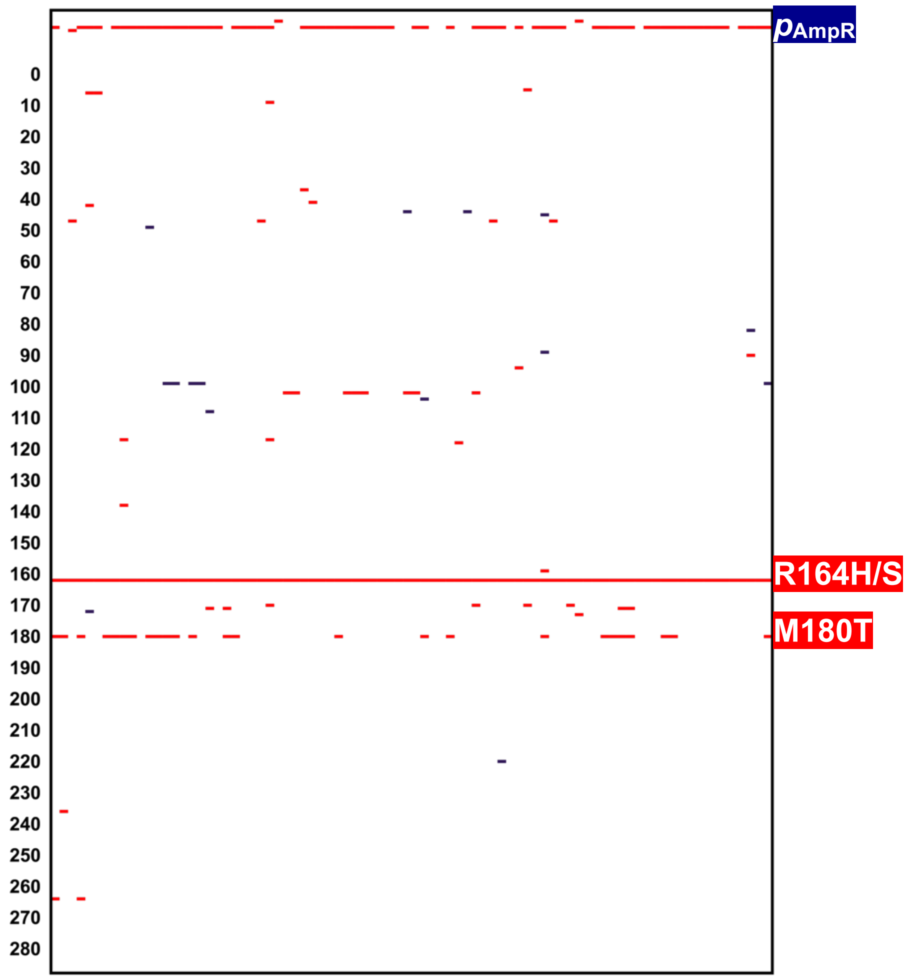


**Figure S13. Sanger sequencing for cefepime resistance continuous evolution experiment.** Convergence of sequences in the promoter region ($p_{\mathrm{AmpR}}$), as well as in the coding sequence (R164H, R164S, and M180T).

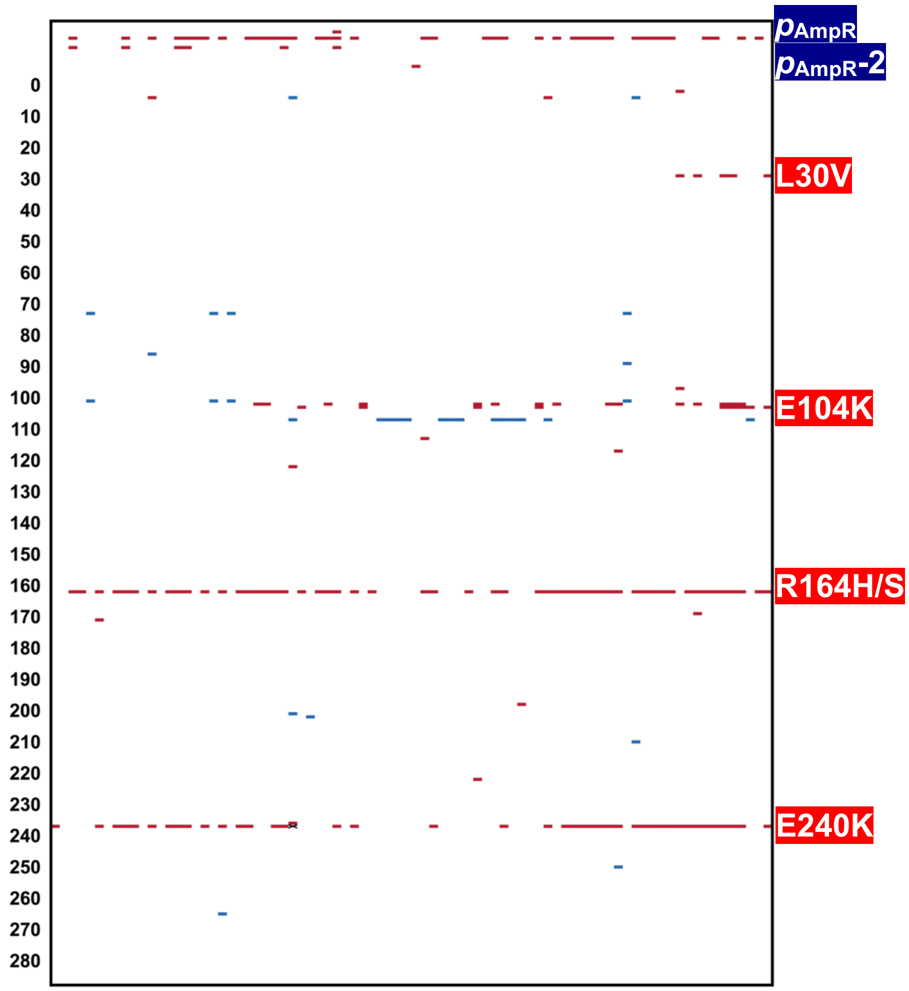


**Figure S14. Sanger sequencing for aztreonam resistance continuous evolution experiment.** Convergence of sequences in the promoter region ($p_{AmpR}$, $p_{AmpR}$-2), as well as in the coding sequence (L30V, E104K, R164H, R164S, and E240K).

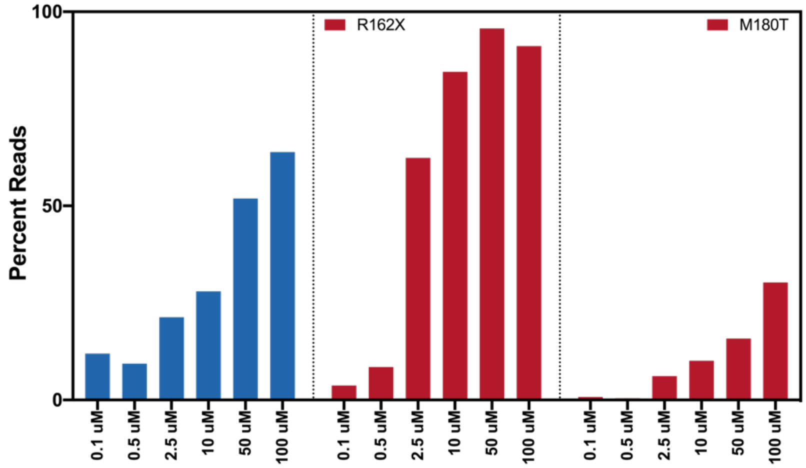


**Figure S15. Time course of β-lactamase evolution experiments.** The time course for evolution of cefepime resistance by deep-sequencing of 96-replicates after each passage (once per day).

Mutations in the promoter are depicted in blue, mutations in the TEM-1 reading frame are depicted in red.

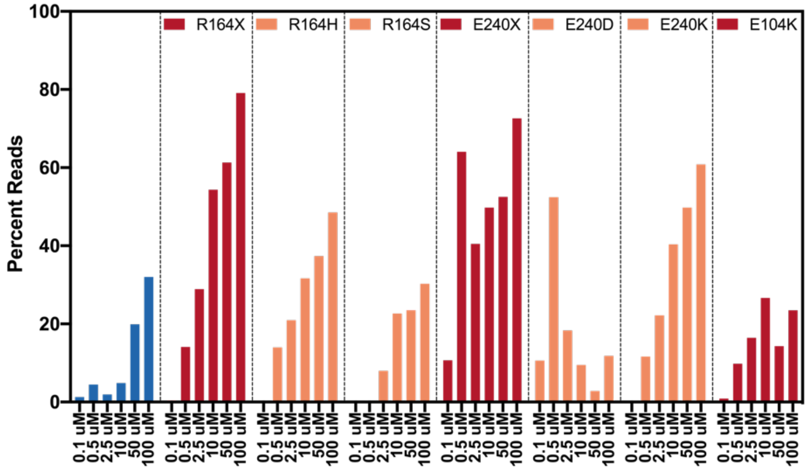


**Figure S16. Time course of β-lactamase evolution experiments.** The time course for evolution of aztreonam resistance was interrogated by deep-sequencing of 96-replicates after each passage (once per day). Mutations in the promoter are depicted in blue, mutations in the TEM-1 reading frame are depicted in red and orange.

### Supplementary Tables

#### Table S1. Plasmids used in this study.

| Name | Specifier | Origin of replication | Gene(s) of interest | Selectable marker |
|---|---|---|---|---|
| **pCD0280** | **pOR1** | pBR322 | $P_{BAD}$::gp4; $P_{GlnS}$::gp2.5 | $Kan^R$ |
| **pC0260** | **pOR2** | pSC101 | $P_{polI}$::gp5 | $Gen^R$ |
| **pCD0261** | **pOR2** | pSC101 | $P_{polI}$::gp5(Δ28) | $Gen^R$ |
| **pCD0262** | **pOR2** | pSC101 | $P_{polI}$::gp5(Δ28, N520M) | $Gen^R$ |
| **pCD0263** | **pOR2** | pSC101 | $P_{polI}$::gp5(Δ28, N520M, P560V) | $Gen^R$ |
| **pCD0264** | **pOR2** | pSC101 | $P_{polI}$::gp5(Δ28, N520M, P560V, V443K) | $Gen^R$ |
| **pCD0265** | **pOR2** | pSC101 | $P_{polI}$::gp5(Δ28, N520M, P560V, V443K, Y24F) | $Gen^R$ |
| **pCD0266** | **pOR2** | pSC101 | $P_{polI}$::gp5(Δ28, N520M, P560V, V443K, Y24F, Q585R) | $Gen^R$ |
| **pCD0275** | **pOR3** | CloDF3 | $P_{proD}$::gp3.5–gp1 | $Sm^R$ |
| **pOR-1** | **pOR-1** | *φ*OR (*7*) | *N.A.* | $Cam^R$ |
| **pOR-2** | **pOR-2** | *φ*OR[1]; pR6K[2] | *N.A.* | $Amp^R$ |
| **pOR-3** | **pOR-3** | *φ*OR | $P_{AmpR}$::*bla*$_{TEM-1}$(Oc) | $Cam^R$ |
| **pOR-4** | **pOR-4** | *φ*OR | $P_{AmpR}$:: *bla*$_{TEM-1}$ | $Cam^R$ |
| **pSR-1** | **pSR-1** | pUC | *dxs* | $Amp^R$ |

#### Table S2. Strains used in this study.

| Strain | Base Strain | Replisome Plasmids | Selectable marker |
|---|---|---|---|
| **T7** $_{\textbf{Rep-Δgp5}}$ | BW25113 | pCD0280, pCD0275 | $Kan^R$, $Sm^R$ |
| **T7**$_{\textbf{Rep-1}}$ | BW25113 | pCD0280, pCD0260, pCD0275 | $Kan^R$, $Gen^R$, $Sm^R$ |
| **T7**$_{\textbf{Rep-2}}$ | BW25113 | pCD0280, pCD0261, pCD0275 | $Kan^R$, $Gen^R$, $Sm^R$ |
| **T7**$_{\textbf{Rep-3}}$ | BW25113 | pCD0280, pCD0262, pCD0275 | $Kan^R$, $Gen^R$, $Sm^R$ |
| **T7**$_{\textbf{Rep-4}}$ | BW25113 | pCD0280, pCD0263, pCD0275 | $Kan^R$, $Gen^R$, $Sm^R$ |

| | | | |
|---|---|---|---|
| **$T7_{Rep-5}$** | BW25113 | pCD0280, pCD0264, pCD0275 | *Kan*$^R$, *Gen*$^R$, *Sm*$^R$ |
| **$T7_{Rep-6}$** | BW25113 | pCD0280, pCD0265, pCD0275 | *Kan*$^R$, *Gen*$^R$, *Sm*$^R$ |
| **$T7_{Rep-7}$** | BW25113 | pCD0280, pCD0266, pCD0275 | *Kan*$^R$, *Gen*$^R$, *Sm*$^R$ |

**Table S3. Mutations rates on orthogonal T7 replicon.**

| Strain | T7 ori Plasmid | T7 DNAP Mutations | Mutation Rate [spb] | Upper 95% CI [spb] | Lower 95% CI [spb] | Copy Number |
|---|---|---|---|---|---|---|
| **$T7_{Rep-1}$** | pOR-3 | Wild type | $2.8\times10^{-9}$ | $8.0\times10^{-10}$ | $7.0\times10^{-10}$ | 1.5 |
| **$T7_{Rep-2}$** | pOR-3 | Δ28 | $3.3\times10^{-8}$ | $9.4\times10^{-9}$ | $8.5\times10^{-9}$ | 12.5 |
| **$T7_{Rep-3}$** | pOR-3 | Δ28, N520M | $6.8\times10^{-7}$ | $1.5\times10^{-8}$ | $1.4\times10^{-8}$ | 5.4 |
| **$T7_{Rep-4}$** | pOR-3 | Δ28, N520M, P560V | $4.3\times10^{-6}$ | $5.5\times10^{-7}$ | $5.2\times10^{-7}$ | 3.8 |
| **$T7_{Rep-5}$** | pOR-3 | Δ28, N520M, P560V, R443K | $1.0\times10^{-5}$ | $1.6\times10^{-6}$ | $1.5\times10^{-6}$ | 4.6 |
| **$T7_{Rep-6}$** | pOR-3 | Δ28, N520M, P560V, R443K, Y24F | $1.1\times10^{-5}$ | $1.5\times10^{-6}$ | $1.5\times10^{-6}$ | 5.0 |
| **$T7_{Rep-7}$** | pOR-3 | Δ28, N520M, P560V, R443K, Y24F, Q585R | $1.7\times10^{-9}$ | $2.6\times10^{-10}$ | $2.5\times10^{-10}$ | 4.1 |

**Table S4. Genomic mutation rates.**

| Strain | T7 DNAP Mutations | Mutation Rate [spb] | Upper 95% CI [spb] | Lower 95% CI [spb] |
|---|---|---|---|---|
| **$T7_{Rep-2}$** | Δ28 | $3.6\times10^{-10}$ | $1.6\times10^{-10}$ | $1.4\times10^{-10}$ |
| **$T7_{Rep-7}$** | Δ28, N520M, P560V, R443K, Y24F, Q585R | $3.9\times10^{-10}$ | $1.7\times10^{-10}$ | $1.6\times10^{-10}$ |

**Table S5. Maximum growth rate and doubling time of replisome strains.**

| Strain | T7 ori Plasmid | Max. rate [ΔOD/min] | $T_{double}$ [min] |
|---|---|---|---|
| **$T7_{Rep-1}$** | pOR-3 | $2.34\times10^{-2}$ | 29.7 |
| **$T7_{Rep-2}$** | pOR-3 | $2.46\times10^{-2}$ | 28.2 |
| **$T7_{Rep-3}$** | pOR-3 | $2.11\times10^{-2}$ | 32.9 |
| **$T7_{Rep-4}$** | pOR-3 | $2.37\times10^{-2}$ | 29.3 |
| **$T7_{Rep-5}$** | pOR-3 | $2.17\times10^{-2}$ | 32.0 |
| **$T7_{Rep-6}$** | pOR-3 | $2.45\times10^{-2}$ | 28.3 |
| **$T7_{Rep-6}$** | pOR-3 | $2.36\times10^{-2}$ | 29.4 |

**Table S6. Strain used in TEM-1 β-lactamase evolution experiments.**

| Strain | T7 ori Plasmid | T7 DNAP Mutations | Mutation Rate [spb] | $T_{double}$ [min] | Copy Number | Mutations/(cell× kb×24h) |
|---|---|---|---|---|---|---|
| **T7$_{Rep-7}$** | pOR-4 | Δ28, N520M, P560V, R443K, Y24F, Q585R | $1.72\times10^{-5}$ | 28.3 | 4.1 | 3.59 |

### References


50. T. Baba, T. Ara, M. Hasegawa, Y. Takai, Y. Okumura, M. Baba, K. A. Datsenko, M. Tomita, B. L. Wanner, H. Mori, Construction of Escherichia coli K-12 in-frame, single-gene knockout mutants: the Keio collection. *Mol. Syst. Biol.* **2**, 8–2006 (2006).
51. B. M. Hall, C.-X. Ma, P. Liang, K. K. Singh, Fluctuation AnaLysis CalculatOR: a web tool for the determination of mutation rate using Luria–Delbrück fluctuation analysis. *Bioinformatics* **25**, 1564–1565 (2009).
52. A. H. Badran, D. R. Liu, Development of potent *in vivo* mutagenesis plasmids with broad mutational spectra. *Nat. Commun.* **6**, 1–10 (2015).
53. C. Lee, J. Kim, S. G. Shin, S. Hwang, Absolute and relative QPCR quantification of plasmid copy number in *Escherichia coli*. *J. Biotechnol.* **123**, 273–280 (2006).
54. R. P. Ambler, A. F. Coulson, J.-M. Frère, J.-M. Ghuysen, B. Joris, M. Forsman, R. C. Levesque, G. Tiraby, S. G. Waley, A standard numbering scheme for the class A beta-lactamases. *Biochem. J.* **276**, 269 (1991).
